# A prototypical phenotype for neutrophils in lupus: the potential contribution of immune complexes

**DOI:** 10.1101/2025.05.19.653336

**Authors:** Sandrine Huot, Paul R. Fortin, Cynthia Laflamme, Philippe A. Tessier, Martin Pelletier, Marc Pouliot

**Affiliations:** Département de microbiologie-infectiologie et immunologie, Faculté de Médecine, Université Laval.; Centre de Recherche du CHU de Québec-Université Laval, Québec, QC, Canada; Centre de recherche ARThrite, Faculté de Médecine de l’Université Laval, Québec, QC, Canada; Division de rhumatologie, Département de médecine, CHU de Québec-Université Laval, Québec, QC, Canada

**Keywords:** Systemic Lupus Erythematosus, Patients, Cohort, Neutrophils, Inflammation, Immune complexes, Soluble Factors

## Abstract

**Purpose:** Systemic lupus erythematosus is an autoimmune disease hallmarked by a plethora of autoantibodies, interferon-signature, auto-immune complexes, dysregulation of soluble factors in circulation, and abnormal neutrophils. We characterized the neutrophil phenotype in a cohort of lupus patients and assessed the implication of the cells environment.

**Methods:** Blood samples were analyzed for neutrophil expression of surface markers and viability, by flow cytometry. Neutrophils from healthy volunteers were stimulated with serum samples from lupus patients. Plasma samples were subjected to multiplex analysis. Whole blood and isolated neutrophils were stimulated with lupus-relevant soluble factors or with heat-aggregated IgGs that mimic the engagement of Fc gamma (Fcγ) receptors by immune complexes, for viability and activation analysis.

**Results:** Lupus neutrophils displayed significant alterations in the expression of surface markers of adhesion, complement regulation, degranulation, and immune complex response. They also had reduced viability and increased apoptosis. Stimulation of healthy neutrophils with lupus serum increased apoptosis. Lupus-relevant soluble factors accelerated neutrophil apoptosis. Heat-aggregated IgGs mirrored most of the key alterations in viability and surface marker expression observed in lupus neutrophils.

**Conclusion:** This study offers a prototypical phenotype for neutrophils in lupus, characterized by shortened viability, partial degranulation, heightened adhesion capacity, and responsiveness to complement activation. It also points to Fcγ receptors engagement as a major driver of the phenotype observed herein. Finally, results emphasize the therapeutic potential of targeting neutrophil-immune complex interactions and the inflammatory plasma milieu in lupus, from which neutrophil dysfunction is largely acquired.

**HIGHLIGHTS:**

- Neutrophils from patients with lupus exhibit abnormal phenotype and viability.
- Analysis of plasmas in lupus reveals altered levels of analytes.
- Stimulation of healthy whole blood with a model of immune complexes reproduces most of the neutrophil phenotypic abnormalities observed in lupus.
- Neutrophil abnormalities in lupus are largely acquired, rather than representing subsets.

**GRAPHICAL ABSTRACT:** 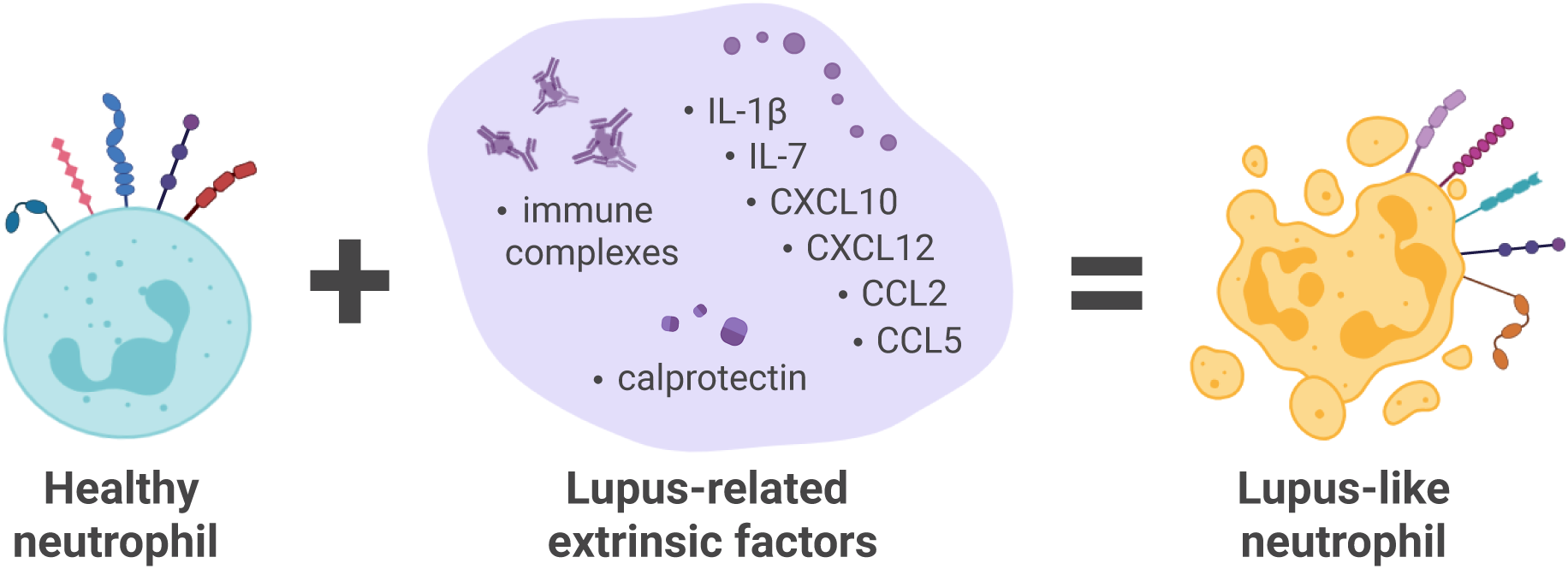

## INTRODUCTION

Systemic lupus erythematosus (SLE) is a complex autoimmune disease characterized by inflammation and immune-mediated injury to multiple organ systems [1, 2]. Approximately 3.4 million people worldwide have been diagnosed with SLE, 90% of whom are women, predominantly of reproductive age [3].

Lupus involves both environmental and genetic factors, innate and adaptive immune activation, and a breakdown of self-tolerance leading to the production of autoantibodies against endogenous antigens, immune complex formation, and subsequent tissue injury [4]. Hallmarks of lupus include an interferon signature [5], a plethora of autoantibodies [6, 7], high levels of immune complexes [7], dysregulation of soluble factors in circulation [8], and neutrophils with altered phenotypes [9]. Environmental triggers, such as infections and ultraviolet light exposure, also exacerbate disease activity by promoting immune activation [10]. Several processes can affect immune cell activation that contributes to cycles of chronic inflammation, tissue injury, and immune dysregulation. The interferon signature, for example, amplifies immune responses and affects monocytes, dendritic cells, and neutrophils [11, 12]. Also, immune complexes trigger inflammation through Fcγ receptors and complement engagement [13, 14].

Neutrophils are the most numerous leukocyte type in circulation and often the first immune cells to migrate to affected tissues and exert host defense functions [15]. In addition to their capacity to engulf and kill microorganisms, neutrophils can engage in intercellular communication and act as influential early regulators that shape subsequent stages of the immune response [16]. These short-lived cells are endowed with remarkable phenotypic plasticity to fulfill their many specialized functions [17]. However, inappropriate neutrophil activation can lead to complications in several autoimmune and inflammatory conditions, including lupus [18]. In fact, neutrophils are key players in lupus pathogenesis, contributing to immune activation and tissue damage [19, 20]. Several potential biomarkers for lupus are neutrophil-related, including granular proteins and modified histones resulting from NET formation. [21]. NET-derived proteins and enzymes not only serve as a source of autoantigens, but also negatively affect vascular health by inducing endothelial apoptosis and oxidizing lipoproteins [22]. Low-density neutrophils (LDNs), enriched in SLE, contribute to excessive type I interferon production [20], which, along with enhanced neutrophil production of tumor necrosis factor (TNF), B-cell stimulating factors like B-cell activating factor (BAFF), and a proliferation-inducing ligand (APRIL), can drive B and T cell abnormalities [23, 24]. Lupus neutrophils also display epigenetic modifications and genomic alterations that may be relevant to their deleterious roles in SLE [21]. Finally, apoptotic neutrophils are more abundant in lupus, which can contribute to tolerance failure, autoimmune lymphocyte activation, and tissue damage. Dying neutrophils may also progress to secondary necrosis, further amplifying autoantigen exposure and immune complex formation, exacerbating the disease [25, 26].

In this study, we sought to characterize the phenotype of circulating neutrophils in a cohort of lupus patients, by analyzing a collection of their surface markers. Our work identifies a prototypical lupus profile of neutrophils and suggests that factors engaging Fcγ receptors contribute in a major way to this profile.

## MATERIALS AND METHODS

### Material

Dextran-500 was purchased from Sigma-Aldrich (Oakville, ON, Canada). Lymphocyte separation medium was purchased from Wisent (St-Bruno, QC, Canada). Lyophilized human immunoglobulins G (IgGs), purified from human plasma or serum by fractionation (purity of greater than 97% as determined by SDS-PAGE analysis) were purchased from Innovative Research, Inc (Novi, MI, USA). Pharm Lyse™ lysing solution and FITC Annexin V Apoptosis Detection Kit were purchased from BD Biosciences (San Diego CA, USA). ProcartaPlex Human Cytokine & Chemokine Panel 1A 34-plex was purchased from Thermo Fisher Scientific (Burlington, ON, Canada). RealTime-Glo™ assay was purchased from Promega (Madison, WI, USA). Recombinant Human IL-1β was purchased from InvivoGen (San Diego, CA, USA). Recombinant Human CXCL12/SDF-1, Recombinant Human CCL5/RANTES, and Recombinant Human CCL2/MCP-1 were purchased from PreproTech (Cranbury, NJ, USA). Recombinant Human IL-7 was purchased from R&D Systems (Minneapolis, MN, USA). Recombinant Human CXCL10/IP-10 and Recombinant Human S100A8/A9 heterodimer (carrier-free), also named calprotectin, were purchased from BioLegend (San Diego, CA, USA). Recombinant Human CXCL1/GRO-α was purchased from Cedarlane (Burlington, ON, Canada).

### Antibodies

APC-labeled mouse anti-human CD11b^act^ (CBRM1/5) was purchased from BioLegend (San Diego, CA, USA). This CBRM1/5 antibody recognizes an activated form of human CD11b. PercP-Cy5.5™-labeled mouse anti-human CD15 (Clone HI98 also known as HIM1), PercP-Cy5.5™-labeled mouse anti-human CD16 (3G8), PE-labeled mouse anti-human CD32 (Clone FLI8.26, also known as 8.26), BV421-labeled mouse anti-human CD35 (E11), APC-labeled mouse anti-human CD46 (E4.3), FITC-labeled mouse anti-human CD55 (IA10), PE-labeled mouse anti-human CD59 (Clone 282 (H19)), V450-labeled mouse anti-human CD64 (10.1), FITC-labeled mouse anti-human CD66b (G10F5), APC-labeled mouse anti-human CD184 (12G5), PE-labeled mouse anti-human CD63 (H5C6), V450-labeled mouse anti-human CD62L (DREG-56), and FITC mouse anti-human CD93 (R139) were purchased from BD Biosciences (San Jose, CA, USA). A list of antibodies used, along with their fluorochromes, alternative target names, and other relevant details, is presented in **Table 1**.

**Table 1.** Details of antibodies used.

| **Target** | **Other name** | **Conjugate** | **Clone** | **Catalog #** | **Monitoring** | **Company** |
| --- | --- | --- | --- | --- | --- | --- |
| CD11b<sup>act</sup> | CBRM | APC | CBRM1/5 | 301410 | Adhesion | Biolegend |
| CD15 | Lewis X (Le<sup>x</sup>) antigen | PercP-Cy5.5 | HI98 | 560828 | Granulocytes, adhesion | BD Biosciences |
| CD16 | FcγRIII | PercP-Cy5.5 | 3G8 | 560717 | Neutrophils, interaction with IgG, secretory vesicles | BD Biosciences |
| CD32 | FcγRII | PE | FLI8.26 | 550586 | Interaction with IgG | BD Biosciences |
| CD35 | CR1/C3bR | BV421 | E11 | 565327 | Complement, secretory vesicles | BD Biosciences |
| CD46 | MCP | APC | E4.3 | 564253 | Complement | BD Biosciences |
| CD55 | DAF | FITC | IA10 | 555693 | Complement | BD Biosciences |
| CD59 | MIRL , MAC-inhibitory protein | PE | p282(H19) | 555764 | Complement | BD Biosciences |
| CD62L | L-selectin | V450 | DREG-56 | 560440 | Adhesion | BD Biosciences |
| CD63 | LAMP-3 | PE | H5C6 | 556020 | Azurophilic (primary) granules | BD Biosciences |
| CD64 | FcγRI | V450 | 10.1 | 561202 | Interaction with IgG | BD Biosciences |
| CD66b | CEACAM8, CGM6, NCA-95 | FITC | G10F5 | 555724 | Specific (secondary) granules, adhesion | BD Biosciences |
| CD93 | C1qR1 | FITC | R139 | 551531 | Complement, apoptosis, secretory vesicles | BD Biosciences |
| CD184 | CXCR4 | APC | 12G5 | 560936 | Chemotaxis, ageing | BD Biosciences |

<sup>act</sup>: Activated form, DAF: Decay-Accelerating Factor, MCP: Membrane Cofactor Protein, MIRL: Membrane Inhibitor of Reactive Lysis, MAC: Membrane Attack Complex.

### Ethics

All experiments involving human tissues received approval from the research ethics committee of CHU de Québec-Université Laval (2022-6235).

### SLE cohort

Participants were diagnosed according to the 1997 Updated American College of Rheumatology (ACR) Classification Criteria [27, 28] and provided consent as required. Blood samples were collected by peripheral vein puncture on isocitrate anticoagulant solution and used within two hours following collection, during their routine visit in the clinic. All clinical data were obtained from the biobank and database of systemic autoimmune rheumatic diseases (SARD) at CHU de Québec-Université Laval. [27, 28]

### Clinical and laboratory variables

Demographic data, including age, biological sex and self-identified ethnicity, as well as SLE characteristics and medication use, were collected for each visit. Disease duration, organ damage-assessed using the Systemic Lupus International Collaborative Clinics/American College of Rheumatology (SLICC/ACR) Damage Index, a validated measurement of organ damage [29], and disease activity-assessed by the SLE Disease Activity Index – 2000 (SLEDAI-2K), a validated lupus activity measure [30], were recorded. Clinical laboratory results (e.g., complete blood counts, urinalysis, complement levels, anti-double-stranded DNA antibodies) were used to calculate SLEDAI-2K scores. Biological samples were paired with demographics and clinical data collected on the corresponding dates.

### Blood collection from healthy volunteers

Informed consent was obtained in writing from all donors. Data collection and analyses were performed anonymously. Blood samples from healthy subjects were collected by peripheral vein puncture on isocitrate anticoagulant solution.

### Multiplex analysis

Thirty four analytes were measured in plasma from healthy volunteers and individuals with lupus using Luminex bead technology and ELISA assays. Luminex measurements were performed using the ProcartaPlex Human Cytokine & Chemokine Panel 1A 34-plex, following the manufacturer’s instructions. The assays were run on a Bio-Plex 100 system (Bio-Rad, Hercules, CA, USA), and data acquisition was conducted using Bio-Plex Manager Software, Version 6.0. Plasma levels of calprotectin were analyzed by sandwich ELISA as described [31]. Analytes exhibiting the most significant alterations were identified as “lupus factors” for subsequent experiments. The summary of analyte concentrations used in the experimental cocktails and those measured in plasma is provided in **Table 2**.

**Table 2.** Analyte concentrations in plasma and experimental cocktails.

| Analyte | Plasma concentration<br>(pg/mL) |  | Experimental cocktail concentration<br>(pg/mL) |  |  |  |
| --- | --- | --- | --- | --- | --- | --- |
|  |  |  | Isolated cells |  | Whole blood |  |
|  | Ctrl | SLE | Ctrl | SLE | Ctrl | SLE |
| IL-1β | 0.3 | 5.23 | – | 5 | – | 5 |
| IL-7 | 0.02 | 0.677 | – | 1 | – | 1 |
| CXCL10/IP-10 | 11.04 | 28.79 | 10 | 50 | – | 50 |
| CXCL12/SDF-1 | 416.6 | 740.5 | 400 | 800 | – | 800 |
| CCL5/RANTES | 32.36 | 111.3 | 50 | 100 | – | 100 |
| CCL2/MCP-1 | 23.32 | 71.70 | 50 | 100 | – | 100 |
| Calprotectin | 1546 | 3970 | 1500 | 4000 | – | 4000 |
| CXCL1/GRO-α | 3.32 | 0.74 | 5 | 1 | – | – |

GRO-α : Growth-regulated Oncogene alpha, IP-10: Interferon gamma-induced Protein 10, MCP-1: Monocyte Chemoattractant Protein-1, RANTES: Regulated upon Activation, Normal T cell Expressed and Secreted, SDF-1: Stromal Cell-Derived Factor 1.

### Preparation of heat-aggregated (HA)-IgGs

HA-IgGs were freshly prepared each day as previously described [32] with modifications [33]. Briefly, IgGs were resuspended in Hank’s Balanced Salt Solution (HBSS) at a concentration of 25 mg/mL and heated at 63°C for 75 min to generate aggregates.

### Whole blood stimulations

Venous blood collected from patients with SLE and healthy volunteers was either left untreated and/or treated with (1): “lupus factors” consisting of IL-1β (5 pg/mL), CXCL12/SDF-1 (800 pg/ml), CCL5/RANTES (100 pg/mL), CCL2/MCP-1 (100 pg/mL), CXCL10/IP-10 (50 pg/mL), IL-7 (1 pg/mL), and calprotectin (4 ng/mL), or with (2): HA-IgGs (1 mg/mL, or as indicated) for 30 min at 37°C. After stimulation, venous blood (100 μl) was incubated with Pharm Lyse™ Buffer (1.9 mL) for 20 min at RT in the dark. After centrifugation (200 x *g*, 5 min), white blood cells were washed with 2 mL of phosphate buffered saline (PBS).

### Viability assay

Neutrophil viability was assessed using a FITC Annexin V Apoptosis Detection Kit, as per the manufacturer’s instructions. Briefly, following appropriate treatments, cell pellets were resuspended in 100 μL of binding buffer, followed by the addition of Annexin V and propidium iodide (PI) to each sample. After 15-min incubation at RT in the dark, 400 μL of binding buffer was added, and samples were analyzed using a FACS Canto II flow cytometer with FACSDiva software, version 6.1.3 (BD Biosciences). Gating was established using control samples labeled individually with Annexin V or PI. Annexin V/PI double-positive cells were identified as late apoptotic/necrotic neutrophils.

### Cell staining

Following appropriate treatments, cell pellets were resuspended in HBSS containing 10 mM HEPES, 1.6 mM Ca^2+^ and no Mg^2+^ and distributed into three separate tubes per condition (100 μL/tube), each containing a specific antibody panel. The first tube contained APC-labeled mouse anti-human CD46, BV421-labeled mouse anti-human CD35, FITC-labeled mouse anti-human CD55, and PE-labeled mouse anti-human CD59; the second contained APC-labeled mouse anti-human CD184, FITC-labeled mouse anti-human CD66b, and PercP-Cy5.5™-labeled mouse anti-human CD15; and the third contained PercP-Cy5.5™-labeled mouse anti-human CD16, V450-labeled mouse anti-human CD64, PE-labeled mouse anti-human CD32, and APC-labeled mouse anti-human CD11b^act^. Cells were incubated for 30 min at 4°C in the dark, then washed and resuspended in 400 μL of 1% paraformaldehyde before flow cytometry analysis. Acquisition was performed with a FACS Canto II flow cytometer with FACSDiva software, version 6.1.3 (BD Biosciences). Neutrophils were identified based on granulocyte gating.

### Purified neutrophils experiments

Neutrophils were isolated as originally described [34] with modifications [35]. Briefly, venous blood collected from healthy volunteers was centrifuged (250 x *g*, 10 min), and the resulting platelet-rich plasma was discarded. Leukocytes were obtained following sedimentation of erythrocytes in 2% Dextran-500. Neutrophils were then separated from other leukocytes by centrifugation on a 10 mL lymphocyte separation medium. Contaminating erythrocytes were removed using 20 sec of hypotonic lysis. Purified granulocytes (> 95% neutrophils, < 5% eosinophils) contained less than 0.2% monocytes, as determined by esterase staining. Viability was greater than 98%, as determined by trypan blue dye exclusion. The entire cell isolation procedure was carried out under sterile conditions at room temperature.

For cytokine-stimulated experiments, two distinct cocktails were prepared. The first cocktail mirrored the “lupus factors” used for whole blood experiments (see *Whole blood stimulations* section), with the addition of CXCL1/GRO-α (1 pg/mL final concentration). The second cocktail represented healthy/normal conditions and included the following: CXCL12/SDF-1 (400 pg/mL), CCL5/RANTES (50 pg/mL), CCL2/MCP-1 (50 pg/mL), CXCL10/IP-10 (10 pg/mL), calprotectin (1.5 ng/mL), and CXCL1/GRO-α (5 pg/mL). The concentrations of the factors in both cocktails, as well as those measured in plasma samples, are summarized in **Table 2**.

### Real-time apoptosis assay

Neutrophil-enriched cell suspensions (0.5x10^6^ cells/condition) were incubated with 3% of autologous, healthy heterologous or lupus serum. Phosphatidylserine (PS) exposure was assessed using the RealTime-Glo™ Annexin V Assay (Promega). Suspensions were incubated at 37°C in the microplate reader Infinite M1000 PRO with i-control 2.0 software (Tecan, Morrisville, NC, USA). Luminescence intensity was monitored every 15 min for 15 hours.

### Statistical analysis

All analyses were performed using GraphPad Prism version 10 (GraphPad Software, San Diego, CA, USA). Where applicable, data are presented as median ± interquartile range. Specific statistical tests are indicated in the corresponding figure legends. Significance thresholds are denoted as follows: *P < 0.05, **P < 0.01, ****P < 0.0001.

## RESULTS

### Study population

**Table 3** summarizes demographic characteristics, lupus disease characteristics and medications. In total, the 98 lupus patients were 86% female with a mean age (SD) of 50.0 (15.9) years and a mean disease duration of 9.5 (10.8) years, and most had low to moderate disease activity mean (SD) SLEDAI = 2.7 (3.3). Biological samples were available from 66 healthy controls. Their mean age was 44.4 year and 71.2% were females. The study population was almost exclusively Caucasian.

**Table 3.** Characteristics of the healthy control group and SLE cohort.

|  | **Total** |  | **Viability cohort** |  | **Surface marker cohort** |  | **Serum cohort** |  | **Plasma analyte cohort** |  |
| --- | --- | --- | --- | --- | --- | --- | --- | --- | --- | --- |
| **Demographics** | Ctrl (n=66) | SLE (n=98) | Ctrl (n=11) | SLE (n=12) | Ctrl (n=25) | SLE (n=43) | Ctrl (n=10) | SLE (n=40) | Ctrl (n=30) | SLE (n=30) |
| ***Age*** |  |  |  |  |  |  |  |  |  |  |
| Range, years | 23-67 | 18-77 | 25-57 | 21-69 | 23-60 | 18-75 | 40-66 | 23-78 | 39-67 | 22-77 |
| Mean ± SD, years | 44.4 ± 11.0 | 50.0 ± 15.9 | 39.3 ± 11.8 | 49.9 ± 16.1 | 40.1 ± 11.5 | 47.6 ± 14.9 | 50.3 ± 7.5 | 53.4 ± 17.0 | 50.2 ± 7.7 | 49.9 ± 16.8 |
| ***Sex, n (%)*** |  |  |  |  |  |  |  |  |  |  |
| Male | 19 (28.8) | 12 ((12.2) | 4 (36.4) | 0 (0.0) | 4 (16.0) | 4 (9.3) | 1 (10.0) | 7 (17.5) | 11 (36.7) | 4 (13.3) |
| Female | 47 (71.2) | 86 ((87.8) | 7 (63.6) | 12 (100.0) | 21 (84.0) | 39 (90.7) | 9 (90.0) | 33 (82.5) | 19 (63.3) | 26 (86.7) |
| ***Ethnicity, n (%)*** |  |  |  |  |  |  |  |  |  |  |
| Caucasian | 66 (100.0) | 97 (99.0) | 11 (100.0) | 12 (100.0) | 25 (100.0) | 42 (97.7) | 10 (100.0) | 39 (97.5) | 30 (100.0) | 30 (100.0) |
| Non-caucasian | 0 (0.0) | 1 (1.0) | 0 (0.0) | 0 (100.0) | 0 (0.0) | 1 (2.3) | 0 (0.0) | 1 (2.5) | 0 (0.0) | 0 (0.0) |
| **Clinical features** |  |  |  |  |  |  |  |  |  |  |
| ***Disease duration*** |  |  |  |  |  |  |  |  |  |  |
| Range, years |  | 0-38 |  | 0-34 |  | 0-32 |  | 0-38 |  | 0-34 |
| Mean ± SD, years |  | 9.5 ± 10.8 |  | 11.7 ± 13.2 |  | 9.7 ± 10.1 |  | 8.0 ± 11.0 |  | 7.5 ± 10.5 |
| ***SLEDAI-2K score*** |  |  |  |  |  |  |  |  |  |  |
| Total range |  | 0-18 |  | 0-6 |  | 0-14 |  | 0-18 |  | 0-18 |
| Mean ± SD |  | 2.7 ± 3.3 |  | 1.9 ± 2.1 |  | 1.7 ± 2.5 |  | 3.7 ± 4.0 |  | 4.5 ± 4.4 |
| ***SLICC DI*** |  |  |  |  |  |  |  |  |  |  |
| Total range |  | 0-13 |  | 0-3 |  | 0-13 |  | 0-9 |  | 0-2 |
| Mean ± SD |  | 0.9 ± 2.2 |  | 0.3 ± 0.9 |  | 1.2 ± 3.0 |  | 0.9 ± 1.7 |  | 0.5 ± 1.0 |
| ***Medication, n (%)*** |  |  |  |  |  |  |  |  |  |  |
| Antimalarial |  | 74 (75.5) |  | 10 (83.3) |  | 33 (76.7) |  | 28 (70.0) |  | 22 (73.3) |
| NSAID |  | 22 (22.4) |  | 6 (50.0) |  | 3 (7.0) |  | 8 (20.0) |  | 9 (30.0) |
| Steroids |  | 21 (21.4) |  | 4 (33.3) |  | 7 (16.3) |  | 11 (27.5) |  | 7 (23.3) |
| Biologics |  | 6 (6.1) |  | 1 (8.3) |  | 3 (7.0) |  | 1 (2.5) |  | 2 (6.7) |

NSAID: Nonsteroidal Anti-Inflammatory Drugs, SLEDAI-2K: SLE Disease Activity Index – 2000, SLICC DI: Systemic Lupus International Collaborating Clinics (SLICC) Damage Index.

### Signature expression profile of surface markers on SLE neutrophils

To generate a comprehensive, prototypical profile of neutrophils in lupus, we monitored a large array of surface markers by flow cytometry [33]. Pilot comparative assays between lupus and normal neutrophils in whole blood progressively guided our focus toward 14 surface markers involved in: immune complex recognition; CD16, CD32, CD64, complement response and regulation; CD35, CD46, CD55, CD59, CD93, degranulation; CD66b, CD63, adhesion: CD11b^act^, CD15, CD62L, and the receptor for stromal cell-derived factor-1 CXCL12/SDF-1: CD184. We compared the surface expression of these 14 markers on neutrophils in whole blood samples from 25 healthy volunteers and 43 patients with lupus. Of these markers, 11 showed significant differences in mean fluorescence intensities (MFI) (**Fig. 1A**). Neutrophils from lupus patients exhibited decreased CD16 (FcγRIII) and CD32 (FcγRII), and elevated CD64 (FcγRI). The four complement-related markers - CD35 (CR1), CD46 (MCP), CD55 (DAF), and CD59 (MAC-inhibitory protein) - were significantly upregulated. Markers related to granule mobilization and adhesion, including CD66b (secondary granules) and the adhesion molecule CD11b^act^, also known as complement receptor component 3 (present on secondary/tertiary granules and secretory vesicles), were also increased. Additionally, CD15 (Lewis X antigen), involved in selectin-mediated adhesion, and CD184 (CXCR4), were upregulated on lupus neutrophils. Alongside the increased MFI, the percentage of positive neutrophils increased for CD64, CD46, CD55, CD59, CD11b^act^, and CD184 (Figure 1B). Overall expression trends of these surface markers on lupus neutrophils, relative to healthy volunteers, are illustrated in **Figure 1C** and provide a prototypical signature profile.

**Figure 1.**
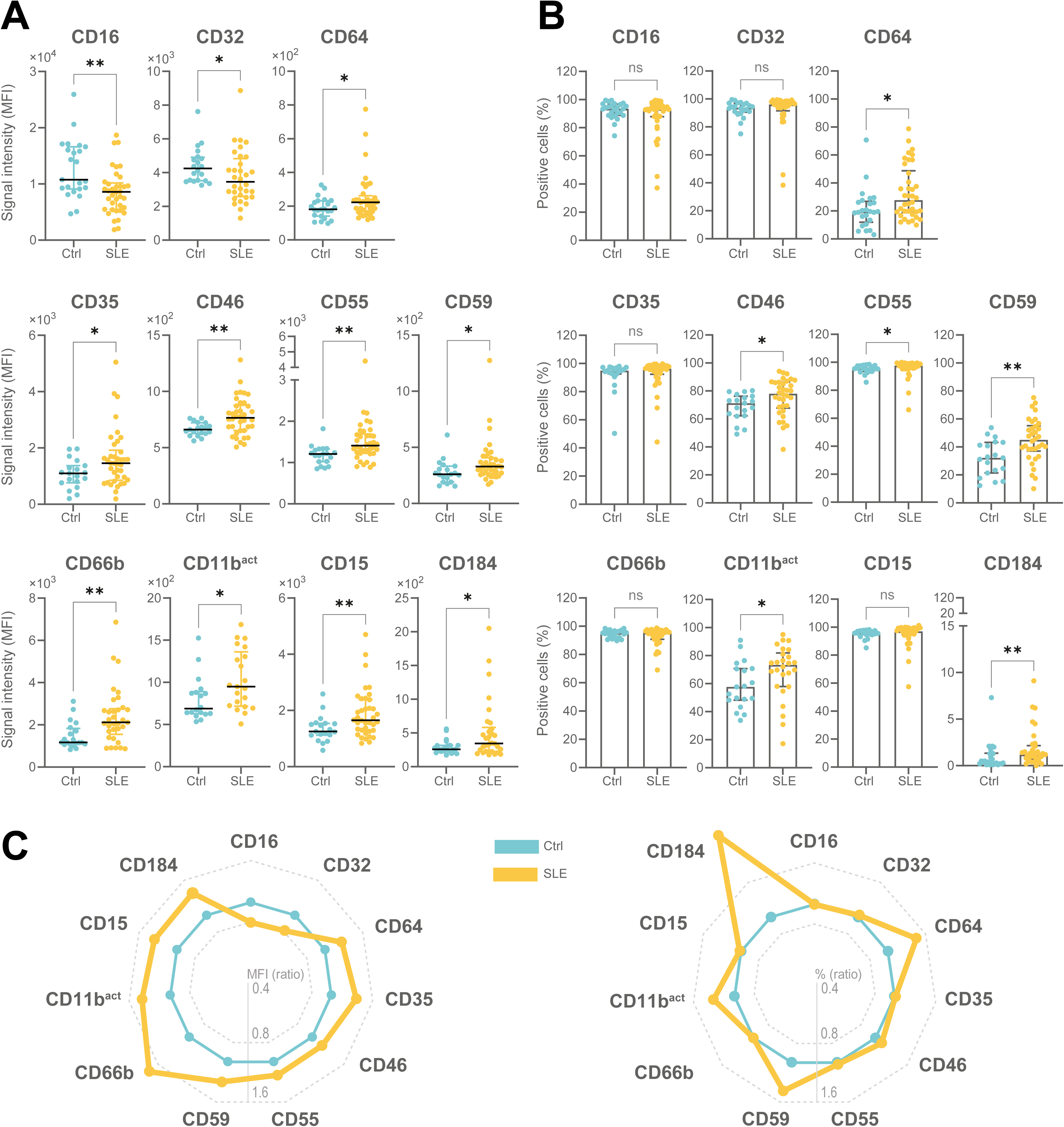
Establishment of a SLE neutrophil phenotypic profile based on surface markers. Whole blood from healthy controls (Ctrl; blue, n = 19–25, depending on the marker) and patients with SLE (orange, n = 21–38, depending on the marker; total cohort = 43) was analyzed by flow cytometry for the indicated surface markers. Neutrophil population was identified through granulocyte gating. Data are shown as median with interquartile range for **(A)** fluorescence signal intensity (MFI) and **(B)** percentage of marker-positive neutrophils. **(C)** Spider chart showing the ratio of median values from SLE patients over healthy donors for MFI (left) and percentage of positive cells (right); values are normalized to the healthy donors. Y-axis is displayed on a log_2_ scale. Statistical analysis was conducted using the Mann-Whitney test. *P < 0.05, **P < 0.01; ns: not significant. ^act^: Activated form, MFI: Mean Fluorescence Intensity. SLE: Systemic Lupus Erythematosus.

### SLE neutrophils show reduced viability and increased apoptosis and sera increased phosphatidylserine exposure

In our cohort, viable neutrophils were lower in patients with lupus than in healthy controls (**Figure 2**), in line with previous reports [25, 36]. This reduction was primarily attributed to a notable twofold increase in early apoptosis in lupus neutrophils. Similar trends were observed after 24 hours, although differences lost statistical significance, and disappeared at 48 hours. To determine whether this alteration resulted from inherently defective neutrophils or initiated by external factors, we incubated neutrophils from a healthy donor with serum samples obtained from lupus patients and monitored levels of PS exposure on the surface, used here as a measure of apoptosis. Compared with autologous or heterologous sera from healthy donors, lupus sera caused a pronounced increase in PS externalization, peaking around four hours post-incubation (**Figure 3A**). The higher trend in PS exposure induced by lupus sera was consistently reproduced with neutrophils from three different healthy donors, with overall PS externalization significantly elevated in the presence of lupus sera (**Figure 3B**). Correlations between the three healthy donors confirmed the consistent impact of SLE serum on neutrophil PS exposure (**Figure 3C**), demonstrating that factors present in the serum of lupus patients do affect neutrophils. This prompted us to identify which factor(s) present in the environment could affect the neutrophil surface marker profile in a fashion observed in lupus.

**Figure 2.**
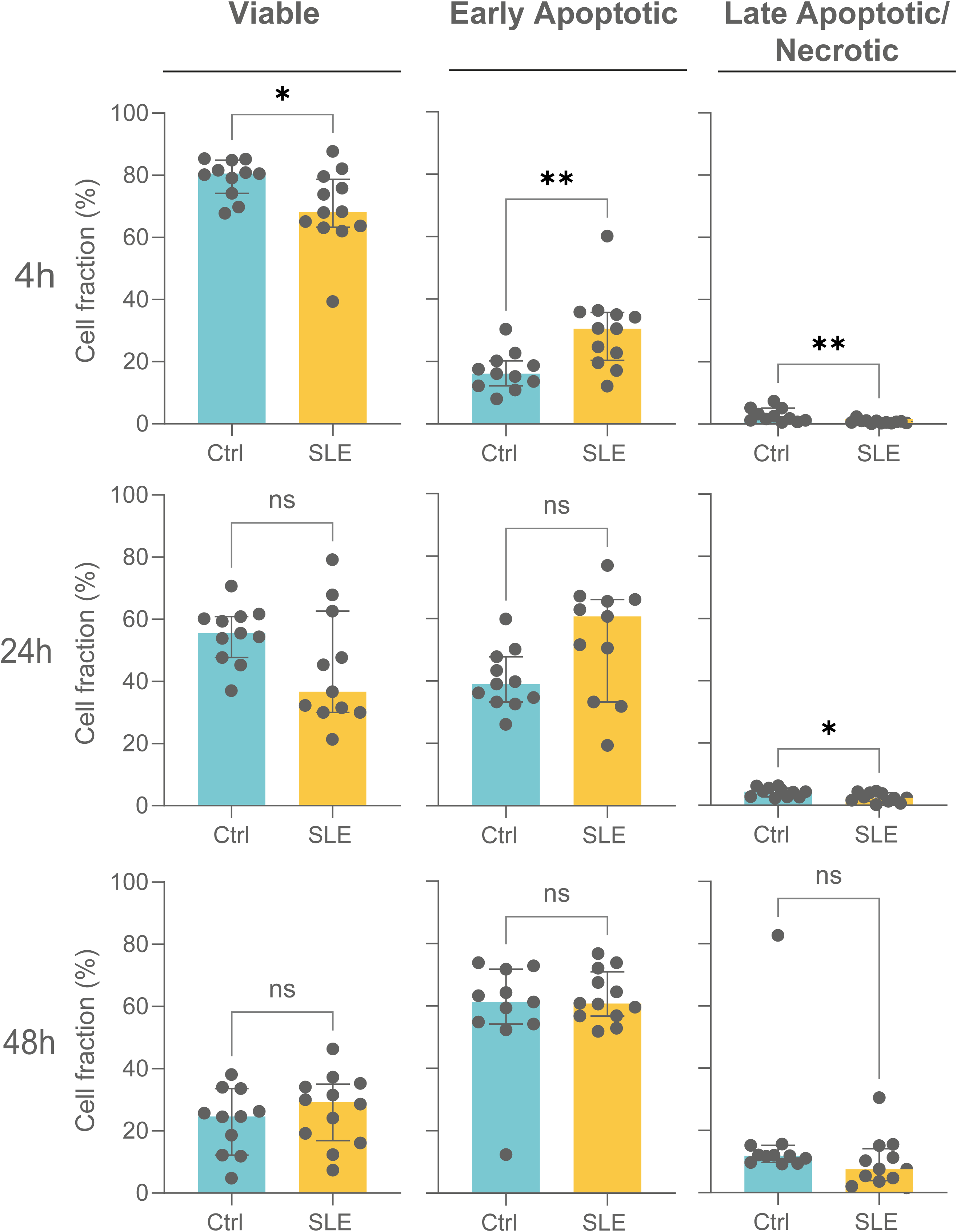
SLE neutrophils exhibit reduced survival and increased apoptosis compared to healthy individuals. Whole blood from healthy controls (Ctrl; blue, n = 11) and patients with SLE (orange, n = 12) were analyzed for apoptosis (FITC-annexin V) and necrosis (propidium iodide) by flow cytometry at the indicated time points (4h, 24h, and 48h). Neutrophil population was identified through granulocyte gating. Data are shown as median with interquartile range. Statistical analysis was performed using the Mann-Whitney test. *P < 0.05, **P < 0.01; ns: not significant. SLE: Systemic Lupus Erythematosus.

**Figure 3.**
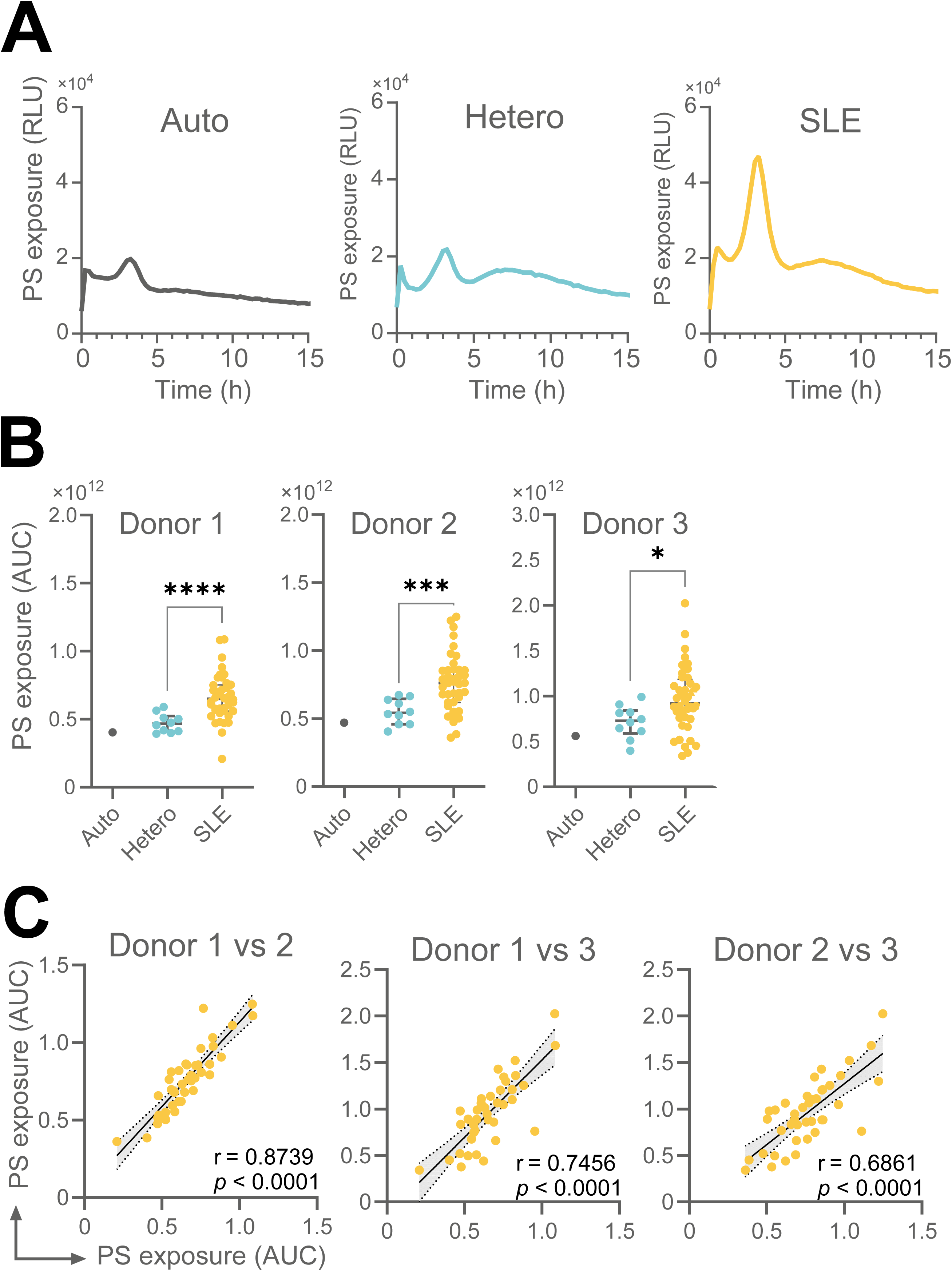
Stimulation of healthy neutrophils with lupus serum increases phosphatidylserine exposure at 4h. Neutrophil-enriched cell suspensions, freshly isolated from healthy volunteers, were incubated with autologous serum (Auto), heterologous serum from healthy donors (Hetero), or serum from individuals with lupus (SLE). **(A)** Representative individual line graphs showing real-time phosphatidylserine (PS) exposure. Data are expressed as relative luminescence units (RLU). **(B)** Results from neutrophil-enriched cell suspensions of three independent donors incubated with their autologous serum, 10 heterologous sera, or 40 SLE sera. **(C)** Inter-donor correlations of PS exposure in response to SLE sera. Data are presented as area under the curve (AUC) derived from the RLU graphs. Statistical analysis was performed using the Kruskal-Wallis test followed by Dunn’s multiple comparisons or the Spearman’s rank correlation test. *P < 0.05, **P < 0.01, ***P < 0.001. SLE: Systemic Lupus Erythematosus.

### Multiplex analysis of analytes in lupus

We employed the ProcartaPlex Human Cytokine & Chemokine Panel 1A 34-plex platform to simultaneously measure over thirty interleukins, chemokines, growth factors, and myeloid-related proteins in plasma samples from SLE patients and healthy controls. We chose plasma samples over serums, for these analyses, based on a previous report indicating improved sensitivity for detecting low-abundance cytokines [37]. Of the analytes measured, seven were markedly elevated (P < 0.0001) in lupus plasma: IL-1β, IL-7, CXCL10/IP-10, CXCL12/SDF-1, CCL2/MCP-1, CCL5/RANTES, and calprotectin (**Figure 4A**). These factors will be referred to hereafter as the *lupus factors* because of the salient differences, in terms of fold-increases, compared to the control group. IL-1RA was also elevated, though to a lesser extent (**Figure 4B**). Conversely, IL-18 and CXCL1/GRO-α were significantly reduced in lupus plasmas. The remaining 24 analytes did not differ significantly between groups. Notably, IFN-α levels were similarly low in both SLE and control samples.

**Figure 4.**
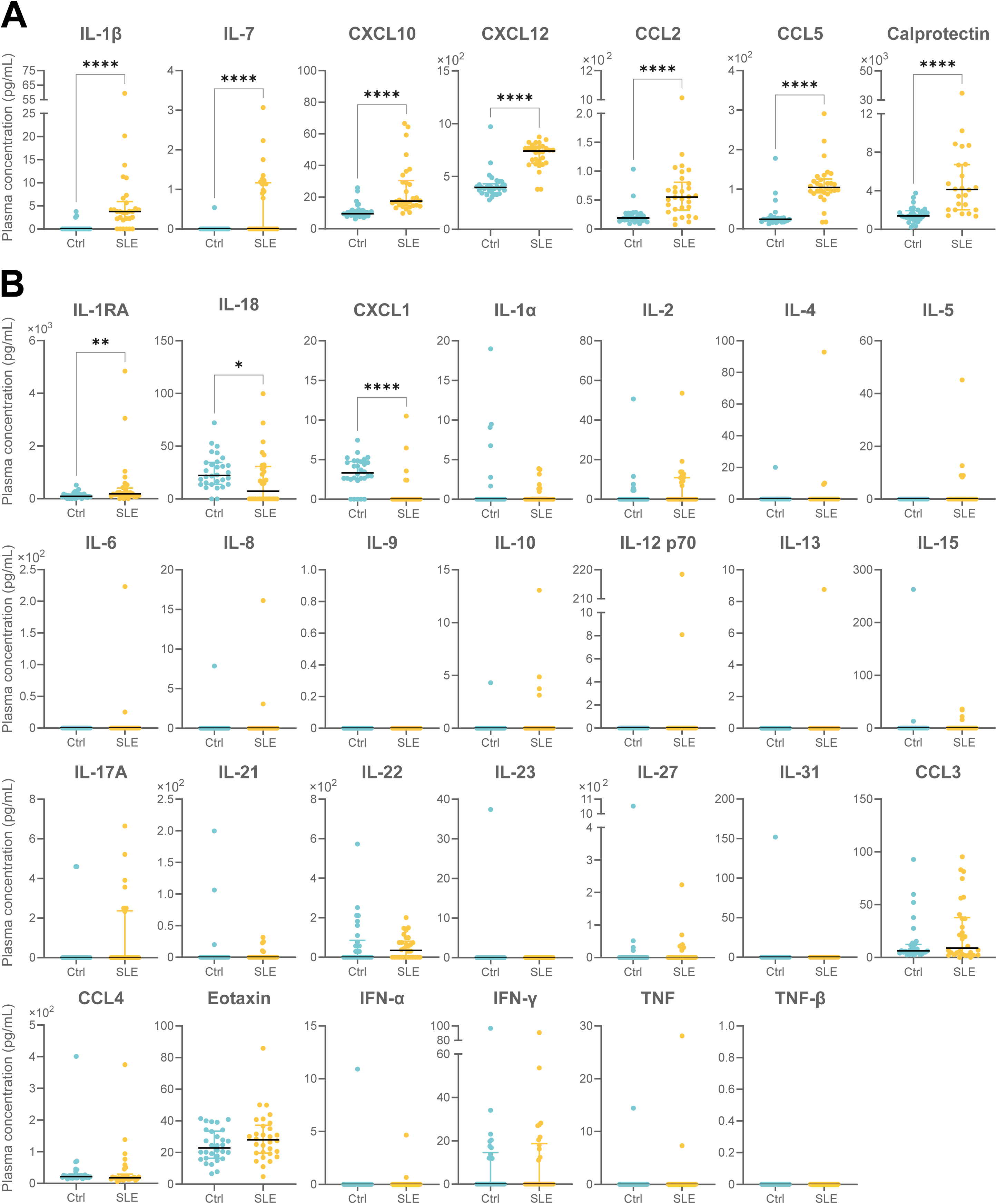
Establishment of SLE plasma analyte profile. Plasma from healthy controls (Ctrl; blue, n = 30) and patients with SLE (orange, n = 30) were analyzed using multiplex assays to quantify over 30 analytes. **(A)** Analytes significantly elevated in SLE plasma identified as lupus-relevant soluble factors (IL-1β, IL-7, CXCL10/IP-10, CXCL12/SDF-1α, CCL2/MCP-1, CCL5/RANTES, and calprotectin). **(B)** Additional analytes, including those with modest increases or no significant differences between groups. Results are shown as median with interquartile range. Statistical analysis was performed using the Mann-Whitney test. *P < 0.05, **P < 0.01, ****P < 0.0001. SLE: Systemic Lupus Erythematosus.

### *Lupus factors* have an impact on neutrophils

Exposure of whole blood from healthy volunteers to *lupus factors* identified above significantly reduced neutrophil viability and increased early apoptosis (**Figure 5**). Also, these factors significantly increased the surface expression of CD35, CD59, and CD66b on neutrophils (**Figure 6A**), as well as the percentage of CD59-positive cells (**Figure 6B**). However, the expression of other surface markers remained unchanged.

**Figure 5.**
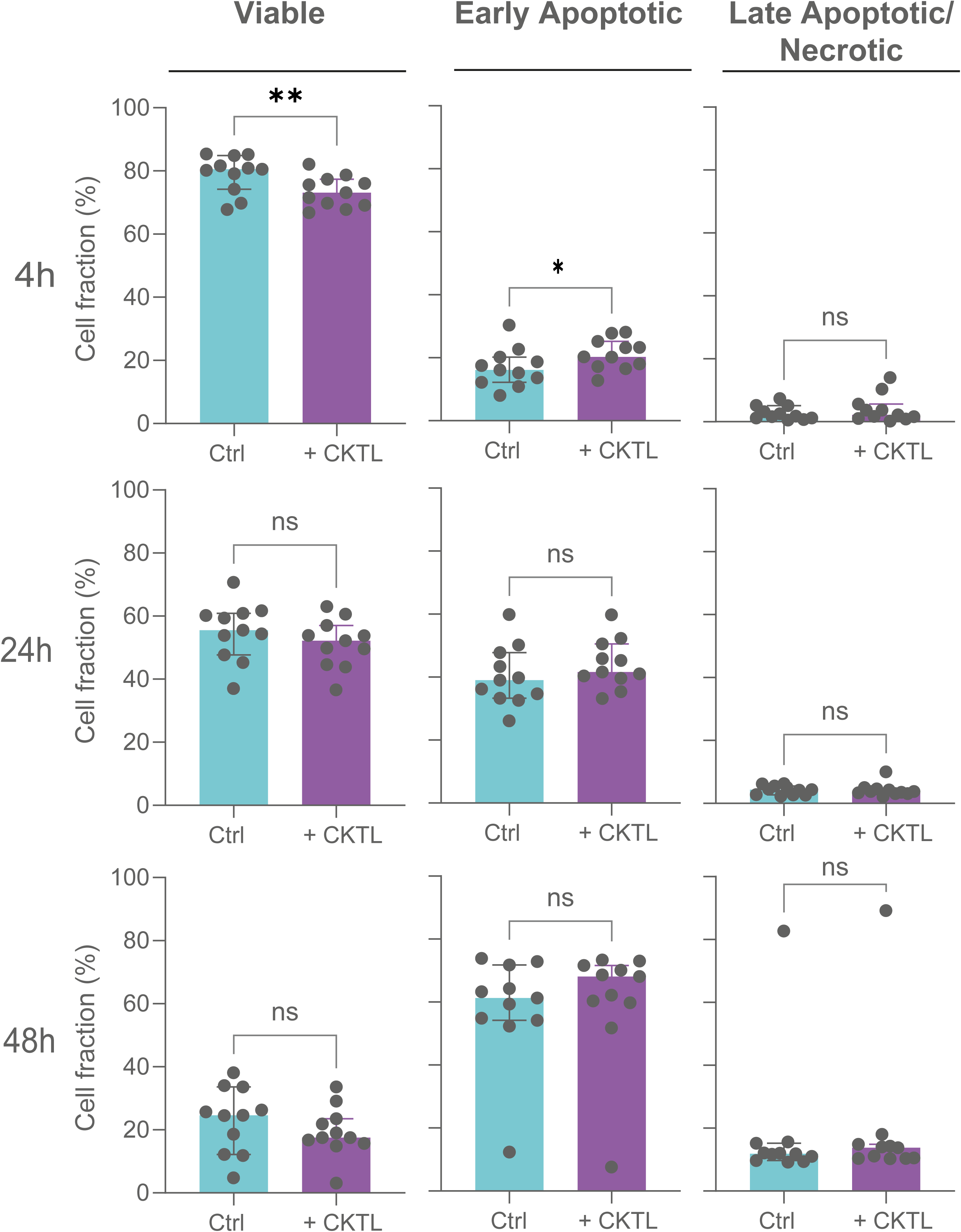
Neutrophils stimulated with the lupus-relevant soluble factors cocktail show reduced survival and increased apoptosis compared to resting neutrophils. Whole blood from healthy volunteers (n=11) was either left untreated (Ctrl; blue) or stimulated with the lupus-relevant soluble factors cocktail (+CKTL; purple) at 37°C for the indicated time points (4h, 24h, and 48h). Neutrophil apoptosis (FITC-annexin V) and necrosis (propidium iodide) were measured by flow cytometry, and neutrophils were identified through granulocyte gating. Statistical comparisons were performed using the Wilcoxon matched pairs signed rank test. *P < 0.05, **P < 0.01, ns: not significant. CKTL: Cocktail.

**Figure 6.**
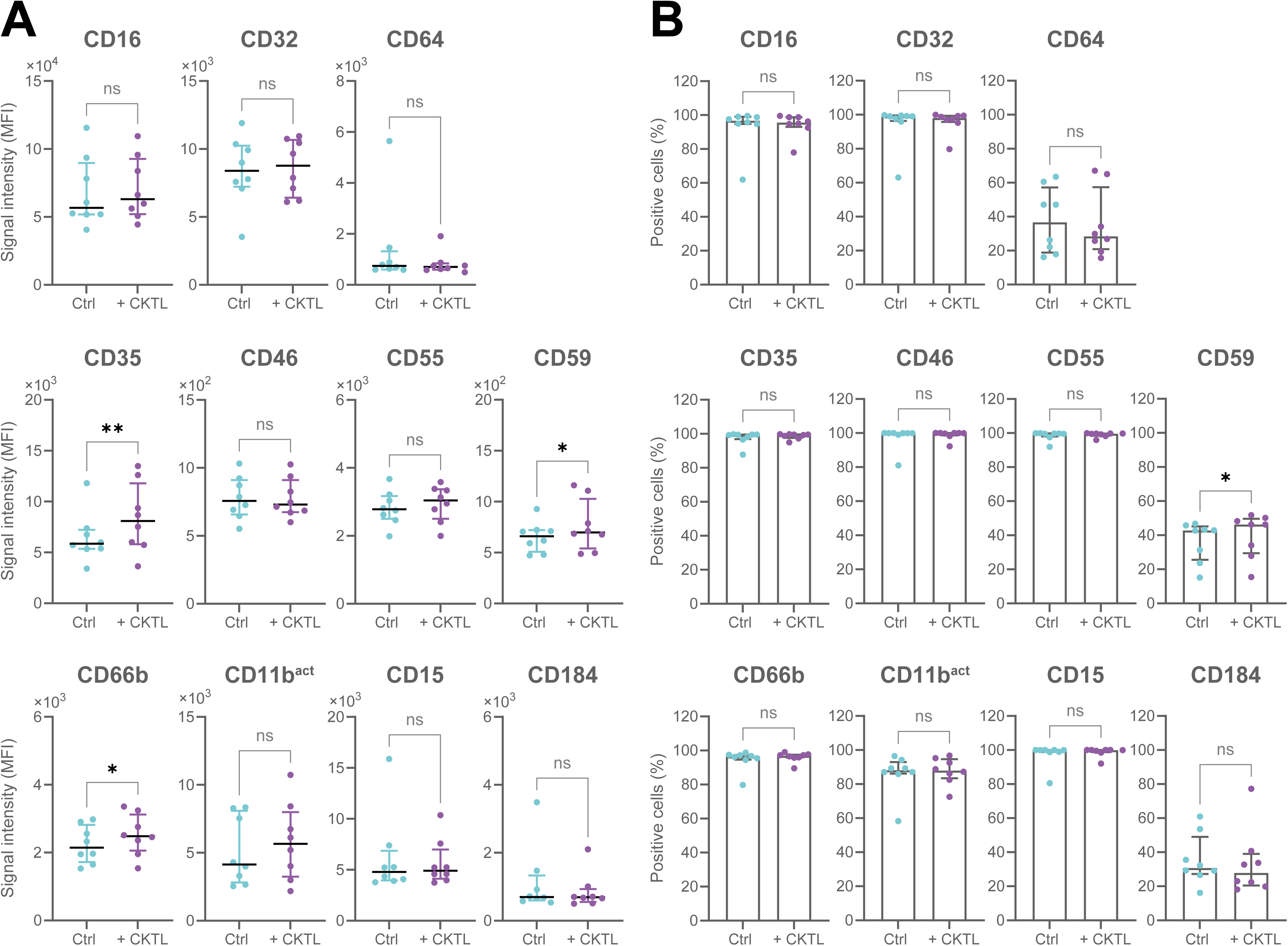
Stimulation with lupus-relevant soluble factors alters the expression of a limited number of surface markers. Whole blood from healthy volunteers (n=8) was either left untreated (Ctrl; blue) or stimulated with the lupus-relevant soluble factors cocktail (+CKTL; purple) at 37°C for 30 min. Surface marker expression was measured by flow cytometry. Neutrophil population was identified through granulocyte gating. Data are shown as median with interquartile range for **(A)** fluorescence signal intensity (MFI) and **(B)** percentage of marker-positive neutrophils. Statistical comparisons were performed using the Wilcoxon matched pairs signed rank test. *P < 0.05, **P < 0.01, ns: not significant. ^act^: Activated form, CKTL: Cocktail, MFI: Mean Fluorescence Intensity.

### Stimulation of whole blood with heat-aggregated-IgGs induces neutrophil phenotypic variations similar to those observed in lupus

Given the important role of immune complexes in lupus pathogenesis, we used heat-aggregated IgGs (HA-IgGs), an antigen-free surrogate for immune complexes, engaging Fcγ receptors [38], and we monitored their impact on neutrophils. In terms of viability, whole blood incubated with HA-IgGs reduced neutrophil viability while increasing the proportion of early apoptotic neutrophils at four hours, in a concentration-dependent manner (**Figure 7)**. Notably, HA-IgGs recapitulated most of the surface marker expression patterns observed in lupus neutrophils. CD32 levels were significantly reduced, while CD64, CD35, CD46, CD55, CD66b, CD11b^act^, and CD15 were significantly upregulated (**Figure 8A**). However, CD59 only showed a modest, non-significant increase; CD16 increased slightly; and CD184 expression remained unchanged. Analysis of the percentage of positive neutrophils also mirrored trends seen in lupus, with increases of several markers, including CD64, CD46, CD59, CD11b^act^, and CD184. However, CD16, CD32, CD66b, and CD15 showed no significant changes (**Figure 8B**).

**Figure 7.**
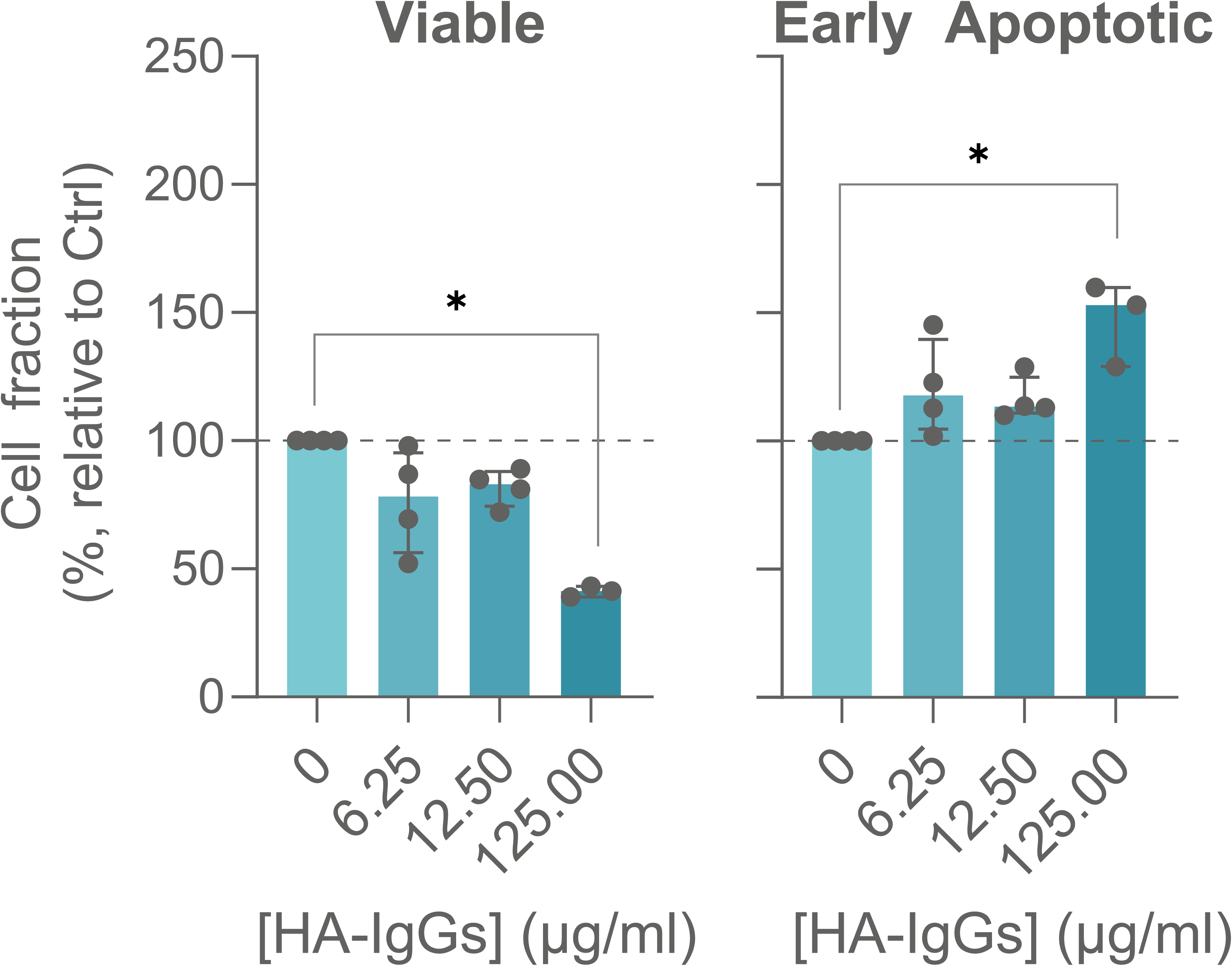
Neutrophil viability and apoptosis exhibit dose-dependent changes in response to HA-IgGs stimulation. Whole blood from healthy controls (n=3-4) was incubated at 37°C with increasing concentrations of HA-IgGs for four hours and analyzed for apoptosis (FITC-annexin V) by flow cytometry. Neutrophil population was identified through granulocyte gating. Cell percentages are normalized to control (Ctrl; 0 µg/ml HA-IgGs) for comparison. Statistical analysis was performed using the Friedman test followed by Dunn’s multiple comparisons. *P < 0.05. HA: Heat-Aggregated.

**Figure 8.**
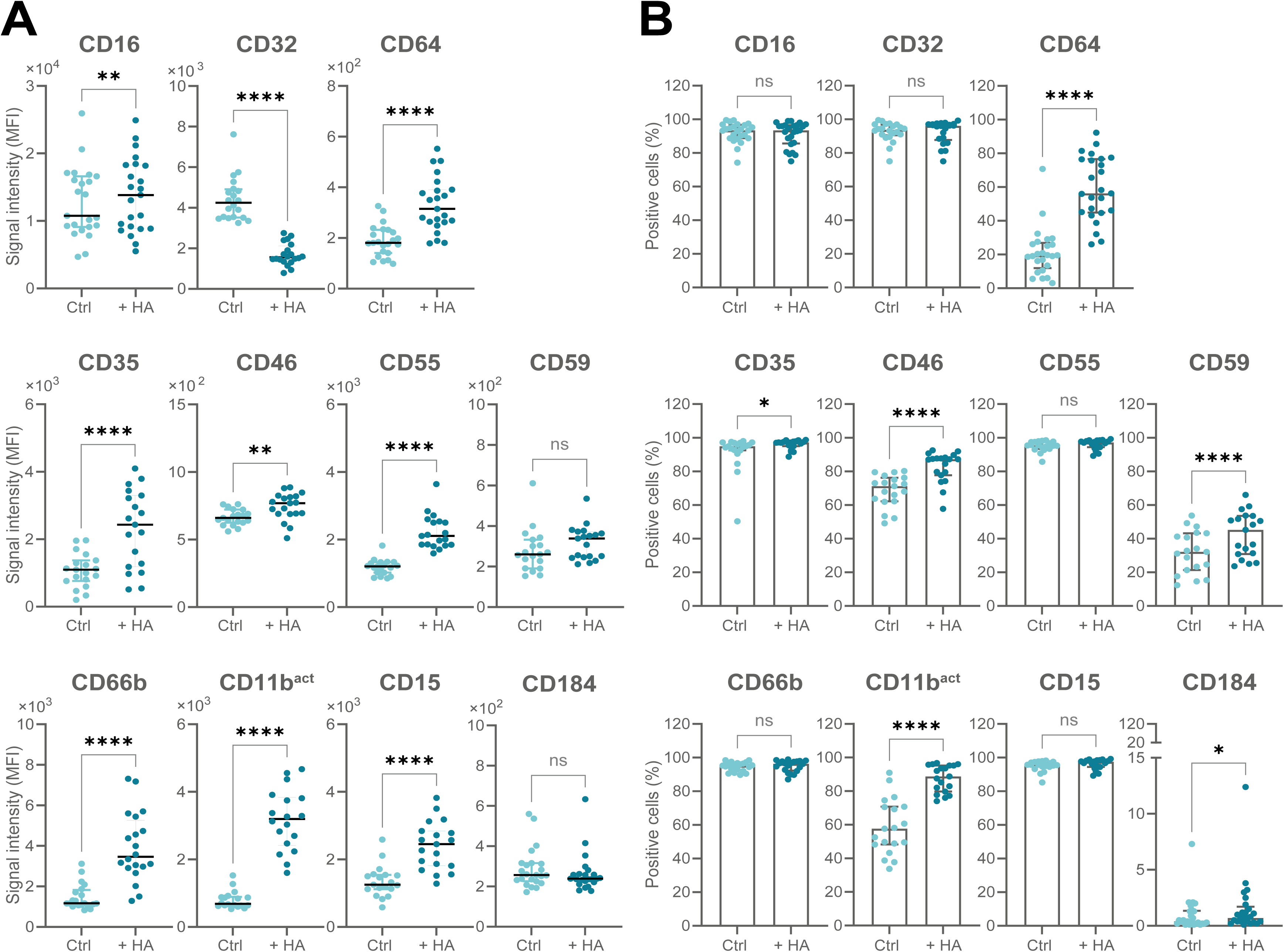
Stimulation of whole blood with HA-IgGs strongly alters the neutrophil phenotypic profile. Whole blood from healthy volunteers (n=19-25) was either left untreated (Ctrl; blue) or incubated with 1mg/ml of HA-IgGs (+HA; dark blue) at 37°C for 30 min. Surface marker expression was measured by flow cytometry and neutrophil population was identified through granulocyte gating. Data are shown as median with interquartile range for **(A)** fluorescence signal intensity (MFI) and **(B)** percentage of marker-positive neutrophils. Statistical comparisons were performed using the Wilcoxon matched pairs signed rank test. *P < 0.05, **P < 0.01, ns: not significant. ^act^: Activated form, HA: Heat-Aggregated, MFI: Mean Fluorescence Intensity.

### HA-IgGs on isolated neutrophils reproduced most of the profile observed in lupus

To further investigate the direct influence of immune complexes on neutrophil phenotype, we performed stimulation assays on isolated neutrophils. HA-IgGs directly caused a significant reduction in CD16 and CD32 expression, alongside significant increases in CD64, CD35, CD55, CD11b^act^, CD15, and CD184 (**Figure 9A**), closely mirroring the alterations observed in lupus whole-blood samples. In terms of percentage of marker-positive cells, HA-IgGs significantly reduced CD32-positive neutrophils while increasing those positive for CD64 and CD184 (**Figure 9B**). A modest increase in CD35-positive cells was also observed. Finally, the frequencies of CD46-, CD55-, CD59-, and CD11b^act^-positive neutrophils were not significantly affected. Taken together, these findings support the notion that Fcγ receptor engagement directly contributes to neutrophil phenotypic alteration.

**Figure 9.**
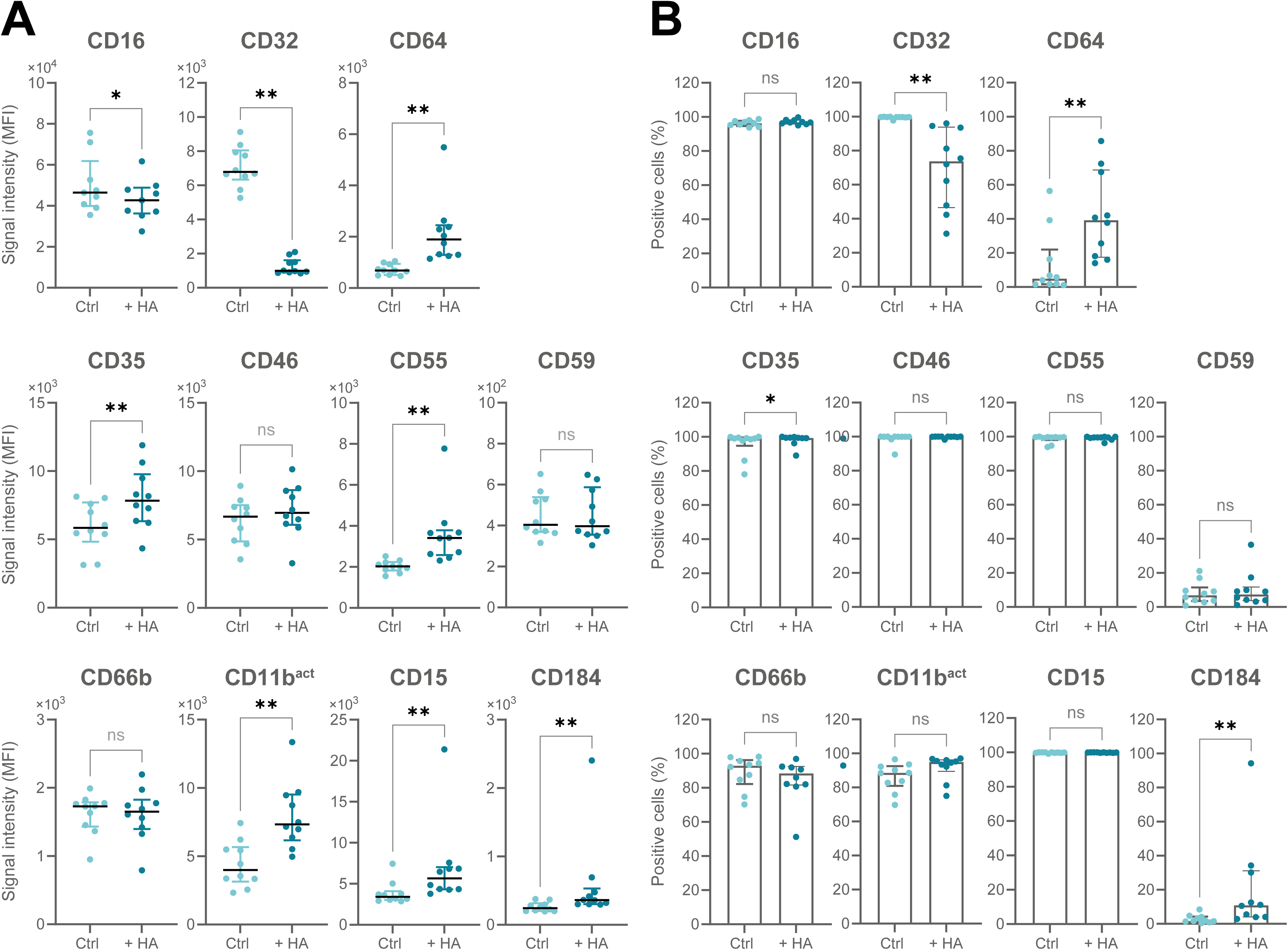
HA-IgGs stimulation alters the neutrophil phenotypic profile in isolated cells. Neutrophil-enriched cell suspensions, freshly isolated from healthy volunteers (n=10), were either left untreated (Ctrl; blue) or incubated with 1mg/ml of HA-IgGs (+HA; dark blue) at 37°C for 30 min. Surface marker expression was measured by flow cytometry. Data are shown as median with interquartile range for **(A)** fluorescence signal intensity (MFI) and **(B)** percentage of marker-positive neutrophils. Statistical comparisons were performed using the Wilcoxon matched pairs signed rank test. *P < 0.05, **P < 0.01, ns: not significant. ^act^: Activated form, HA: Heat-Aggregated, MFI: Mean Fluorescence Intensity.

Finally, to determine whether these phenotypical changes could also occur at concentrations approaching those reported for circulating immune complexes in lupus [39], we conducted concentration-response stimulations with whole blood. MFI increases for CD64, CD55, CD66b, and CD11b^act^ were among the first ones to occur (**Figure 10A**). Regarding percentages of positive cells, CD64 emerged as the most strongly upregulated marker (**Figure 10B**). Together, these findings indicate that HA-IgG stimulation induces several phenotypic hallmarks observed in circulating lupus neutrophils, underscoring the potent influence of immune complexes on neutrophil activation and function.

**Figure 10.**
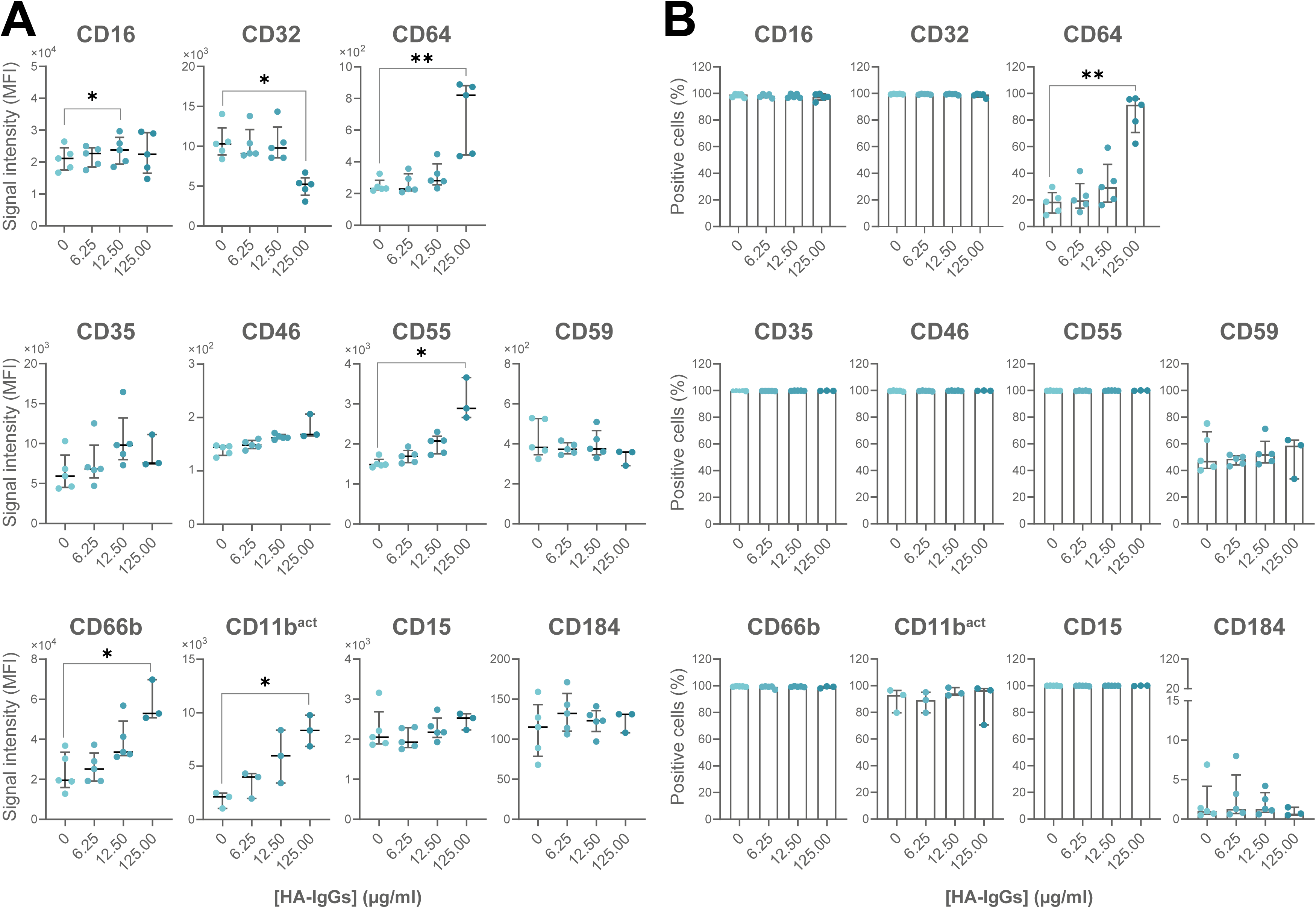
HA-IgGs stimulation induces dose-dependent alterations in the neutrophil phenotypic profile. Whole blood from healthy controls (n=3-5) was incubated with increasing concentrations of HA-IgGs at 37°C for 30 min. Surface marker expression was measured by flow cytometry and neutrophil population was identified through granulocyte gating. Data are shown as median with interquartile range for **(A)** fluorescence signal intensity (MFI) and **(B)** percentage of marker-positive neutrophils. Statistical analysis was performed using the Friedman test followed by Dunn’s multiple comparisons. *P < 0.05, **P < 0.01. ^act^: Activated form, HA: Heat-Aggregated, MFI: Mean Fluorescence Intensity.

## DISCUSSION

This study procures a prototypical signature profile of neutrophil surface markers, in the context of lupus. This profile uncovered new insights, in addition to corroborating previously reported individual trends. Expression of CD16, a low-affinity Fcγ receptor, was reduced, likely reflecting increased apoptosis [25, 40]. Decreased CD32 may result from receptor shedding [41] or internalization following immune complex engagement [42] or even competition between immune complexes and the detection antibody. In contrast, CD64 -the only high-affinity Fcγ receptor- was elevated, suggesting a shift toward heightened immune complex sensitivity and priming for enhanced effector functions such as superoxide production [33, 43]. Complement activation, while critical for immune complex clearance, can propagate tissue injury if dysregulated [44]. Increased complement regulatory proteins (Cregs) on neutrophils may reflect a compensatory mechanism to prevent excessive complement activation [45]. Elevated CD59 expression, along with CD66b, also indicates secondary granule mobilization [46]. Variability in Creg expression in some previous studies [47, 48] may be linked with differences in ethnicity, disease severity or leukocyte subset dynamics, particularly given that neutropenia is observed in a minority of lupus patients [49, 50]. We identified a modest but significant increase in CD184 (CXCR4) expression on circulating lupus neutrophils, consistent with murine models [51]. CD184^hi^ neutrophils may be linked to cellular aging or prior tissue exposure [52] and align with broader patterns of CD184 elevation in lupus B and T cells [53, 54]. Together, these features reinforce a lupus neutrophil phenotype characterized by shortened viability, partial degranulation, heightened adhesion capacity, and responsiveness to complement activation.

Neutrophil plasticity enables functional adaptation across inflammatory contexts [55]. The characteristics of lupus neutrophils unveiled herein appears to be a relevant example of neutrophil plasticity in action [55]. In support, the phenotypic abnormalities, including accelerated apoptosis and specific surface marker expression displayed by lupus neutrophils, were largely induced upon normal neutrophils by lupus-associated extrinsic factors, including immune complex-like particles. Indeed, exposure of healthy neutrophils to HA-IgGs, used as a model of immune complexes engaging Fcγ receptors [38], recapitulated most features observed in lupus neutrophils, suggesting that Fcγ receptor engagement may drive some of the key features found on lupus neutrophils.

Most of the lupus factors that were elevated in the plasma of patients with lupus here have been associated with sustaining chronic inflammation [56-58]. Moreover, calprotectin levels, a marker of neutrophil activation [59], and IL-1β concentrations were also increased, reinforcing the existence of a neutrophil-activating environment. Prior reports on IL-1β levels in lupus have been inconsistent [58, 60], while our findings suggest its modest systemic elevation. IFN-alpha (type I) and IFN-gamma (type II) were barely present, in line with the generally low disease activity levels displayed by our cohort of patients [61]. Despite the clear potential of the soluble factors elevated in lupus within this relatively comprehensive survey, they only reproduced in part the profile observed in lupus neutrophils. Additional factors, yet unidentified, may also be at play.

Acute *in vitro* stimulation by HA-IgGs does not reflect the chronic, low-level stimulation by immune-complexes experienced *in vivo*. Also, HA-IgGs used in this study were prepared from a random mixture of IgGs, likely different from actual ratios and glycosylation levels observed in lupus [62, 63]. Moreover, HA-IgGs do not contain antigens like immune complexes do and likely to produce additional cellular responses, through TLR signaling. Distinct studies should investigate the impact of antigen-specific immune complexes [64] on neutrophil functions. Finally, additional factors distinct ot those monitored herein, are also likely at play in lupus and affecting neutrophils. Nonetheless, it remains remarkable that factors engaging the Fcγ receptors are sufficient to recapitulate much of the lupus neutrophil phenotype observed here.

This prototypical profile of lupus neutrophils is, to our knowledge, the first of its kind. It emanates from a limited number of patients, recruited in our local cohort, and who are in relatively good health and of caucasian ethnicity for the most part. Results are likely to differ with larger and more diverse cohorts of different disease severity. Nonetheless, the present study may represent a starting point for the representation of neutrophils in lupus, and inother autoimmune diseases where immune complexes are involved as well.

In conclusion, by defining key features of lupus neutrophils and identifying candidate mediators of their activation, we provide a framework for future studies aimed at restoring neutrophil homeostasis and mitigating tissue damage in lupus. Our study emphasizes the therapeutic potential of targeting neutrophil–immune complex interactions and the inflammatory plasma milieu in lupus, from which neutrophil dysfunction is largely acquired [47, 65], offering new insights into disease mechanisms and new directions for therapeutic intervention.

## STATEMENTS

### Data availability

Data sharing not applicable to this article as no datasets were generated or analyzed during the current study.

### Ethics approval and consent to participate

All experiments involving human tissues received approval from the research ethics committee of CHU de Québec-Université Laval (2022-6235). All participants provided consent in writing, as required.

### Patient consent statement

All experiments involving human tissues received approval from the research ethics committee of CHU de Québec-Université Laval (2022-6235). Informed consent was obtained in writing from all donors.

### Consent for publication

Clinical data is presented, but on an anonymous basis. All participants provided written consent, as required.

### Permission to reproduce material from other sources

N/A

### Clinical trial registration

N/A

## Acknowledgements

N\A

## Authors contribution

**SH:** Conceptualization, Methodology, Data curation, Writing – original draft, Writing – review and editing.

**PRF:** Conceptualization, Validation.

**CL:** Methodology, Data curation.

**PAT:** Methodology.

**MPe:** Methodology.

**MPo:** Funding acquisition, Conceptualization, Supervision, Data curation, Writing – review and editing, final version of the manuscript.

## REFERENCES

[1] A. Suarez-Fueyo, S. J. Bradley, G. C. Tsokos. T cells in Systemic Lupus Erythematosus. Curr Opin Immunol, 2016;43:32–8.

[2] G. Tsokos. Systemic Lupus Erythematosus: Basic, Applied and Clinical Aspects, 2nd ed. Academic Press; 2020.

[3] G. J. Pons-Estel, G. S. Alarcon, L. Scofield, L. Reinlib, G. S. Cooper. Understanding the epidemiology and progression of systemic lupus erythematosus. Semin Arthritis Rheum, 2010;39:257–68.

[4] C. C. Mok, C. S. Lau. Pathogenesis of systemic lupus erythematosus. J Clin Pathol, 2003;56:481–90.

[5] M. K. Crow. Type I interferon in the pathogenesis of lupus. J Immunol, 2014;192:5459–68.

[6] G. Yaniv, G. Twig, D. B. Shor, A. Furer, Y. Sherer, O. Mozes et al. A volcanic explosion of autoantibodies in systemic lupus erythematosus: a diversity of 180 different antibodies found in SLE patients. Autoimmun Rev, 2015;14:75–9.

[7] M. R. Arbuckle, M. T. McClain, M. V. Rubertone, R. H. Scofield, G. J. Dennis, J. A. James et al. Development of autoantibodies before the clinical onset of systemic lupus erythematosus. N Engl J Med, 2003;349:1526–33.

[8] H. Zhou, B. Li, J. Li, T. Wu, X. Jin, R. Yuan et al. Dysregulated T Cell Activation and Aberrant Cytokine Expression Profile in Systemic Lupus Erythematosus. Mediators Inflamm, 2019;2019:8450947.

[9] L. Bennett, A. K. Palucka, E. Arce, V. Cantrell, J. Borvak, J. Banchereau et al. Interferon and granulopoiesis signatures in systemic lupus erythematosus blood. J Exp Med, 2003;197:711–23.

[10] D. Fernandez, K. A. Kirou. What Causes Lupus Flares? Curr Rheumatol Rep, 2016;18:14.

[11] E. C. Baechler, F. M. Batliwalla, G. Karypis, P. M. Gaffney, W. A. Ortmann, K. J. Espe et al. Interferon-inducible gene expression signature in peripheral blood cells of patients with severe lupus. Proc Natl Acad Sci U S A, 2003;100:2610–5.

[12] J. D. Clough. Role of autoantibodies and immune complexes in the pathogenesis of systemic lupus erythematosus. J Clin Apher, 1992;7:151–2.

[13] S. Ben Mkaddem, M. Benhamou, R. C. Monteiro. Understanding Fc Receptor Involvement in Inflammatory Diseases: From Mechanisms to New Therapeutic Tools. Front Immunol, 2019;10:811.

[14] C. M. Karsten, J. Kohl. The immunoglobulin, IgG Fc receptor and complement triangle in autoimmune diseases. Immunobiology, 2012;217:1067–79.

[15] M. Siwicki, P. Kubes. Neutrophils in host defense, healing, and hypersensitivity: Dynamic cells within a dynamic host. J Allergy Clin Immunol, 2023;151:634–55.

[16] J. Wang, J. Wang. Neutrophils, functions beyond host defense. Cell Immunol, 2022;379:104579.

[17] R. Sumagin. Emerging neutrophil plasticity: Terminally differentiated cells no more. J Leukoc Biol, 2021;109:473–5.

[18] M. Fresneda Alarcon, Z. McLaren, H. L. Wright. Neutrophils in the Pathogenesis of Rheumatoid Arthritis and Systemic Lupus Erythematosus: Same Foe Different M.O. Front Immunol, 2021;12:649693.

[19] M. J. Kaplan. Neutrophils in the pathogenesis and manifestations of SLE. Nat Rev Rheumatol, 2011;7:691–9.

[20] E. Villanueva, S. Yalavarthi, C. C. Berthier, J. B. Hodgin, R. Khandpur, A. M. Lin et al. Netting neutrophils induce endothelial damage, infiltrate tissues, and expose immunostimulatory molecules in systemic lupus erythematosus. J Immunol, 2011;187:538–52.

[21] C. K. Smith, M. J. Kaplan. The role of neutrophils in the pathogenesis of systemic lupus erythematosus. Curr Opin Rheumatol, 2015;27:448–53.

[22] X. Yang, Y. Ma, X. Chen, J. Zhu, W. Xue, K. Ning. Mechanisms of neutrophil extracellular trap in chronic inflammation of endothelium in atherosclerosis. Life Sci, 2023;328:121867.

[23] A. Palanichamy, J. W. Bauer, S. Yalavarthi, N. Meednu, J. Barnard, T. Owen et al. Neutrophil-mediated IFN activation in the bone marrow alters B cell development in human and murine systemic lupus erythematosus. J Immunol, 2014;192:906–18.

[24] H. Zhuang, S. Han, Y. Xu, Y. Li, H. Wang, L. J. Yang et al. Toll-like receptor 7-stimulated tumor necrosis factor alpha causes bone marrow damage in systemic lupus erythematosus. Arthritis Rheumatol, 2014;66:140–51.

[25] Y. Ren, J. Tang, M. Y. Mok, A. W. Chan, A. Wu, C. S. Lau. Increased apoptotic neutrophils and macrophages and impaired macrophage phagocytic clearance of apoptotic neutrophils in systemic lupus erythematosus. Arthritis Rheum, 2003;48:2888–97.

[26] L. E. Munoz, K. Lauber, M. Schiller, A. A. Manfredi, M. Herrmann. The role of defective clearance of apoptotic cells in systemic autoimmunity. Nat Rev Rheumatol, 2010;6:280–9.

[27] E. M. Tan, A. S. Cohen, J. F. Fries, A. T. Masi, D. J. McShane, N. F. Rothfield et al. The 1982 revised criteria for the classification of systemic lupus erythematosus. Arthritis Rheum, 1982;25:1271–7.

[28] M. C. Hochberg. Updating the American College of Rheumatology revised criteria for the classification of systemic lupus erythematosus. Arthritis Rheum, 1997;40:1725.

[29] D. Gladman, E. Ginzler, C. Goldsmith, P. Fortin, M. Liang, M. Urowitz et al. Systemic lupus international collaborative clinics: development of a damage index in systemic lupus erythematosus. J Rheumatol, 1992;19:1820–1.

[30] D. D. Gladman, D. Ibanez, M. B. Urowitz. Systemic lupus erythematosus disease activity index 2000. J Rheumatol, 2002;29:288–91.

[31] M. R. Tardif, J. A. Chapeton-Montes, A. Posvandzic, N. Page, C. Gilbert, P. A. Tessier. Secretion of S100A8, S100A9, and S100A12 by Neutrophils Involves Reactive Oxygen Species and Potassium Efflux. J Immunol Res, 2015;2015:296149.

[32] R. Gale, J. V. Bertouch, T. P. Gordon, J. Bradley, P. J. Roberts-Thomson. Neutrophil activation by immune complexes and the role of rheumatoid factor. Ann Rheum Dis, 1984;43:34–9.

[33] S. Huot, C. Laflamme, P. R. Fortin, E. Boilard, M. Pouliot. IgG-aggregates rapidly upregulate FcgRI expression at the surface of human neutrophils in a FcgRII-dependent fashion: A crucial role for FcgRI in the generation of reactive oxygen species. FASEB J, 2020;34:15208–21.

[34] M. E. Fiset, C. Gilbert, P. E. Poubelle, M. Pouliot. Human neutrophils as a source of nociceptin: a novel link between pain and inflammation. Biochemistry, 2003;42:10498–505.

[35] M. Pouliot, M. E. Fiset, M. Masse, P. H. Naccache, P. Borgeat. Adenosine up-regulates cyclooxygenase-2 in human granulocytes: impact on the balance of eicosanoid generation. J Immunol, 2002;169:5279–86.

[36] C. Y. Tsai, K. J. Li, S. C. Hsieh, H. T. Liao, C. L. Yu. What’s wrong with neutrophils in lupus? Clin Exp Rheumatol, 2019;37:684–93.

[37] Y. Rosenberg-Hasson, L. Hansmann, M. Liedtke, I. Herschmann, H. T. Maecker. Effects of serum and plasma matrices on multiplex immunoassays. Immunol Res, 2014;58:224–33.

[38] K. K. Ostreiko, I. A. Tumanova, K. Sykulev Yu. Production and characterization of heat-aggregated IgG complexes with pre-determined molecular masses: light-scattering study. Immunol Lett, 1987;15:311–6.

[39] A. A. Bengtsson, H. Tyden, C. Lood. Neutrophil FcgammaRIIA availability is associated with disease activity in systemic lupus erythematosus. Arthritis Res Ther, 2020;22:126.

[40] D. A. Moulding, C. A. Hart, S. W. Edwards. Regulation of neutrophil FcgammaRIIIb (CD16) surface expression following delayed apoptosis in response to GM-CSF and sodium butyrate. J Leukoc Biol, 1999;65:875–82.

[41] C. Lood, S. Arve, J. Ledbetter, K. B. Elkon. TLR7/8 activation in neutrophils impairs immune complex phagocytosis through shedding of FcgRIIA. J Exp Med, 2017;214:2103–19.

[42] G. Szucs, M. Kavai, P. Suranyi, E. Kiss, I. Csipo, G. Szegedi. Correlations of monocyte phagocytic receptor expressions with serum immune complex level in systemic lupus erythematosus. Scand J Immunol, 1994;40:481–4.

[43] S. Huot, P. R. Fortin, C. Laflamme, M. Pouliot. Neutrophil FcgammaRI expression as a determinant of oxidative responses in human blood. J Leukoc Biol, 2026;118.

[44] M. Noris, G. Remuzzi. Overview of complement activation and regulation. Semin Nephrol, 2013;33:479–92.

[45] C. Q. Schmidt, J. D. Lambris, D. Ricklin. Protection of host cells by complement regulators. Immunol Rev, 2016;274:152–71.

[46] D. L. Gordon, H. Papazaharoudakis, T. A. Sadlon, A. Arellano, N. Okada. Upregulation of human neutrophil CD59, a regulator of the membrane attack complex of complement, following cell activation. Immunol Cell Biol, 1994;72:222–9.

[47] C. M. Marzocchi-Machado, C. M. Alves, A. E. Azzolini, A. C. Polizello, I. F. Carvalho, Y. M. Lucisano-Valim. Fcgamma and complement receptors: expression, role and co-operation in mediating the oxidative burst and degranulation of neutrophils of Brazilian systemic lupus erythematosus patients. Lupus, 2002;11:240–8.

[48] A. P. Alegretti, L. Schneider, A. K. Piccoli, O. A. Monticielo, P. S. Lora, J. C. Brenol et al. Diminished expression of complement regulatory proteins on peripheral blood cells from systemic lupus erythematosus patients. Clin Dev Immunol, 2012;2012:725684.

[49] L. Carli, C. Tani, S. Vagnani, V. Signorini, M. Mosca. Leukopenia, lymphopenia, and neutropenia in systemic lupus erythematosus: Prevalence and clinical impact--A systematic literature review. Semin Arthritis Rheum, 2015;45:190–4.

[50] N. A. Alhammadi, D. D. Gladman, J. Su, M. B. Urowitz. Isolated Neutropenia in Systemic Lupus Erythematosus. J Rheumatol, 2023;50:459–60.

[51] A. Wang, A. M. Fairhurst, K. Tus, S. Subramanian, Y. Liu, F. Lin et al. CXCR4/CXCL12 hyperexpression plays a pivotal role in the pathogenesis of lupus. J Immunol, 2009;182:4448–58.

[52] S. Skopelja-Gardner, J. Tai, X. Sun, L. Tanaka, J. A. Kuchenbecker, J. M. Snyder et al. Acute skin exposure to ultraviolet light triggers neutrophil-mediated kidney inflammation. Proc Natl Acad Sci U S A, 2021;118.

[53] A. Wang, P. Guilpain, B. F. Chong, S. Chouzenoux, L. Guillevin, Y. Du et al. Dysregulated expression of CXCR4/CXCL12 in subsets of patients with systemic lupus erythematosus. Arthritis Rheum, 2010;62:3436–46.

[54] L. D. Zhao, D. Liang, X. N. Wu, Y. Li, J. W. Niu, C. Zhou et al. Contribution and underlying mechanisms of CXCR4 overexpression in patients with systemic lupus erythematosus. Cell Mol Immunol, 2017;14:842–9.

[55] L. G. Ng, I. Ballesteros, M. A. Cassatella, M. Egeblad, Z. G. Fridlender, D. Gabrilovich et al. From complexity to consensus: A roadmap for neutrophil classification. Immunity, 2025;58:1890–903.

[56] L. C. Lit, C. K. Wong, L. S. Tam, E. K. Li, C. W. Lam. Raised plasma concentration and ex vivo production of inflammatory chemokines in patients with systemic lupus erythematosus. Ann Rheum Dis, 2006;65:209–15.

[57] E. Hrycek, A. Franek, E. Blaszczak, J. Dworak, A. Hrycek. Serum levels of selected chemokines in systemic lupus erythematosus patients. Rheumatol Int, 2013;33:2423–7.

[58] Y. Wu, B. Cai, J. Zhang, B. Shen, Z. Huang, C. Tan et al. IL-1beta and IL-6 Are Highly Expressed in RF+IgE+ Systemic Lupus Erythematous Subtype. J Immunol Res, 2017;2017:5096741.

[59] K. A. Zervides, A. Jern, J. Nystedt, B. Gullstrand, P. C. Nilsson, P. C. Sundgren et al. Serum S100A8/A9 concentrations are associated with neuropsychiatric involvement in systemic lupus erythematosus: a cross-sectional study. BMC Rheumatol, 2022;6:38.

[60] R. Mende, F. B. Vincent, R. Kandane-Rathnayake, R. Koelmeyer, E. Lin, J. Chang et al. Analysis of Serum Interleukin (IL)-1beta and IL-18 in Systemic Lupus Erythematosus. Front Immunol, 2018;9:1250.

[61] K. V. Yerram, R. Baisya, P. Kumar, R. Mylavarapu, L. Rajasekhar. Serum interferon-alpha predicts in-hospital mortality in patients hospitalised with acute severe lupus. Lupus Sci Med, 2023;10.

[62] S. J. Wei, Q. Xiong, H. Yao, Q. M. He, P. L. Yu. Is systemic lupus erythematosus linked to Immunoglobulin G4 Autoantibodies? Hum Immunol, 2024;85:110826.

[63] A. Beyze, C. Larroque, M. Le Quintrec. The role of antibody glycosylation in autoimmune and alloimmune kidney diseases. Nat Rev Nephrol, 2024;20:672–89.

[64] M. Tokuyama, B. M. Gunn, A. Venkataraman, Y. Kong, I. Kang, T. Rakib et al. Antibodies against human endogenous retrovirus K102 envelope activate neutrophils in systemic lupus erythematosus. J Exp Med, 2021;218.

[65] T. Wang, R. Kuley, P. Hermanson, P. Chu, C. Pohlmeyer, J. D. Ravichandar et al. Immune complexes-mediated activation of neutrophils in systemic lupus erythematosus is dependent on RNA recognition by toll-like receptor 8. Front Immunol, 2024;15:1515469.

